# Transcriptomic datasets of melittin- and un-treated murine cervical carcinoma U14 cells

**DOI:** 10.64898/2026.07.25.739748

**Authors:** Ronghua Zhang, Yuwei Zhang, Mengyi Wang, Jianrong Jiang, Yueliang Li, Dafu Chen, Tizhen Yan, Rui Guo

## Abstract

Melittin, a potent amphipathic cationic peptide derived from bee venom, exhibits broad-spectrum antineoplastic efficacy, notably against cervical carcinoma. Despite its established therapeutic potential, the global transcriptional reprogramming orchestrating its acute multi-pathway cytotoxicity remains incompletely understood. To bridge this knowledge gap, we generated the first comprehensive, untargeted RNA-seq dataset profiling the acute phase of melittin-induced cell death in murine cervical carcinoma U14 cells (exposed to 4 μg/mL melittin for 20 minutes) alongside untreated controls. Utilizing deep sequencing and rigorous bioinformatics workflows, we quantified genome-wide mRNA abundances and mapped a distinct transcriptomic shift, identifying 254 significantly differentially expressed genes, comprising 158 up- and 96 down-regulated transcripts. Validated by stringent quality control metrics, exceptional genomic mapping rates, and comprehensive functional annotations via the GO and KEGG databases, this high-resolution transcriptomic resource provides a systems-level map of early molecular alterations. All raw and processed sequencing data are publicly available. This transcriptomic resource provides a valuable foundation for elucidating the acute regulatory networks underlying melittin-induced anti-cervical cancer effects.

**Dataset:** The dataset can be accessed through the NGDC website by searching with the BioProject accession number PRJCA068439. Reviewers may use this link for anonymous access during the review process. Direct URL to data: Genome Sequence Archive-CNCB-NGDC

## 1. Summary

Melittin, a 26-amino-acid amphipathic cationic peptide derived from bee venom, is well-recognized for its potent and broad-spectrum antineoplastic properties^[1]^. By either inserting into biological membranes to compromise their integrity or translocating intracellularly, melittin disrupts diverse signaling cascades^[2,3]^. Accumulating evidence across various human cancer cell lines and rodent models indicates that melittin selectively induces cytotoxicity in malignant cells via multiple cell death modalities, including apoptosis, necrosis, and autophagy^[4-6]^. Particularly in the context of cervical cancer, melittin has exhibited profound therapeutic efficacy. Previous studies have elucidated that melittin potently inhibits the proliferation of cervical carcinoma cells (such as human HeLa and murine U14 models) and triggers mitochondrion-dependent apoptosis by altering the Bax/Bcl-2 expression ratio and downregulating PI3K/Akt signaling pathway^[7]^. However, while scattered studies have documented the modulation of specific transcripts by melittin in certain cancer types^[8]^, the global transcriptional reprogramming orchestrating these multi-pathway cell death events remains incompletely understood.

Evasion of programmed cell death represents a fundamental hallmark of malignancy. Consequently, ideal chemotherapeutic agents must engage multidimensional signaling networks to overcome such resistance^[9]^. Given the pleiotropic nature of melittin, evaluating its systemic efficacy requires going beyond traditional single-gene or isolated-pathway approaches. High-throughput transcriptome sequencing (RNA-seq) provides an unbiased, genome-wide perspective, which is crucial for capturing the dynamic gene expression shifts and intricate regulatory networks triggered by anticancer peptides. To date, a comprehensive RNA-seq dataset dedicated to delineating the global mRNA signatures governed by melittin during the acute phase of cancer cell death is critically lacking. This absence substantially impedes our understanding of melittin’s multi-target mechanisms and the identification of reliable biomarkers.

To bridge this substantial knowledge gap, we performed a comprehensive, untargeted transcriptomic analysis of murine cervical carcinoma U14 cells (*Mus musculus*). Cells were subjected to 4 μg/mL melittin treatment for 20 minutes or left untreated as controls, with five independent biological replicates per condition to ensure statistical stringency.

By utilizing deep sequencing alongside rigorous bioinformatic pipelines, we not only quantified genome-wide mRNA abundances but also mapped a distinct transcriptomic shift, identifying 158 significantly up- and 96 down-regulated genes (screening thresholds: |log_2_(Fold change)| ≥ 1 and *Q* value < 0.05) in response to melittin exposure. To further decode the biological imperatives of these differentially expressed genes (DEGs), systematic functional annotation workflows were then integrated. In strict alignment with our transcriptomic methodology, pathway-level and functional enrichments were meticulously conducted based on the Gene Ontology (GO) and the Kyoto Encyclopedia of Genes and Genomes (KEGG) databases.

The final dataset includes all raw high-throughput sequencing data files (FASTQ format), corresponding MD5 checksum files for data integrity validation, and processed gene expression matrices. All raw data have been deposited in the National Genomics Data Center (NGDC) under BioProject accession number PRJCA068439. The full raw dataset can be accessed directly via the permanent NGDC BioProject link: https://ngdc.cncb.ac.cn/bioproject/browse/PRJCA068439. No login or access permission is required.

This dataset will serve as the core data foundation for our ongoing and future research projects focused on elucidating the transcriptomic-mediated anticancer mechanisms of melittin, and for the preclinical development of multi-pathway therapeutic strategies against cervical carcinoma. To date, no peer-reviewed research articles based on this full dataset have been published.

In conclusion, as the first publicly available transcriptomic resource profiling acute melittin exposure in murine cervical carcinoma cells, this dataset delivers a comprehensive, systems-level map of early global gene expression alterations. By delineating these acute transcriptional mechanisms, it effectively bridges the knowledge gap left by conventional long-term or targeted signaling studies. Ultimately, this high-resolution RNA-seq repository establishes an essential cornerstone for future integrative multi-omics analyses, empowering the global scientific community to decode the novel mechanisms of membrane-active peptides and accelerate the translational development of natural bioactive peptide-based, broad-spectrum antineoplastic therapeutics.

## 2. Data Description

### 2.1 Dataset Structure

This dataset is deposited in the NGDC (National Genomics Data Center) Genome Sequence Archive under BioProject accession number PRJCA068439. It contains only raw sequencing data files organized as described below. No processed data or additional documentation folders are included in this submission; only the raw data files are provided.

### 2.2 Raw Data Folder Structure

Raw RNA sequencing data files are organized by experimental group and deposited in the Genome Sequence Archive (Genomics, Proteomics & Bioinformatics 2025) in National Genomics Data Center (Nucleic Acids Res 2026), China National Center for Bioinformation /Beijing Institute of Genomics, Chinese Academy of Sciences (GSA: CRA046053). All data are publicly accessible at https://ngdc.cncb.ac.cn/gsa. The files for each sample are listed as follows:

## 3. Methods

### 3.1 Cell culture and melittin treatment

Murine cervical carcinoma U14 cells (obtained from Immocell Biotechnology, Xiamen, China) were cultured in high-glucose Dulbecco’s Modified Eagle Medium (DMEM, Gibco, USA) supplemented with 10% fetal bovine serum (FBS, Gibco, USA) and 1% penicillin-streptomycin (100 U/mL penicillin and 100 μg/mL streptomycin) at 37°C in a humidified incubator containing 5% CO_2_. Logarithmic-phase cells were trypsinized and seeded into 6-well plates at a density of 2 × 10^^6^ cells per well. For morphological evaluation, sterile glass coverslips were pre-inserted into the wells prior to seeding. Following a 24 hours incubation to allow for complete cell attachment and acclimation, U14 cells were divided into a melittin-treated (T) group and an un-treated control (C) group. Melittin (purity ≥ 98%, Selleck, USA) was dissolved in complete culture medium to prepare a stock solution. The treatment group was exposed to a final concentration of 4 μg/mL melittin for exactly 20 min, a condition optimized based on prior viability assays. Five independent biological replicates (n = 5) were prepared per condition to ensure rigorous statistical stringency.

Upon treatment completion, the samples were parallelly processed. Cells designated for morphological analysis were directly observed and imaged using an inverted phase-contrast microscope (Sunny Optical, China) without fixation to evaluate melittin-induced morphological alterations (Figure 1). Concurrently, cells designated for transcriptome profiling (C1-C5, T1-T5) were immediately harvested, snap-frozen in liquid nitrogen, and subjected to total RNA extraction.

**Figure 1.**
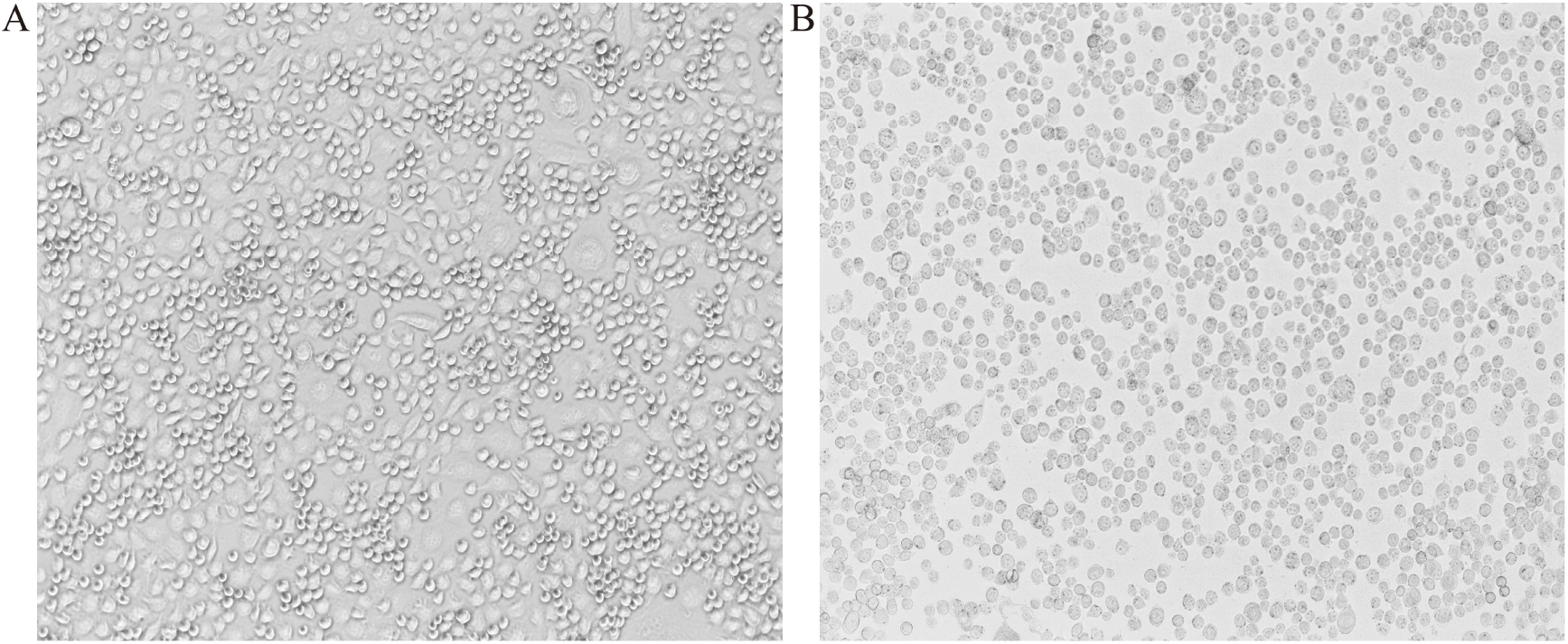
Inverted optical microscopy observation of melittin-induced morphological alterations in U14 cells (10X magnification). (A) Un-treated control group, (B) Melittin-treatment group at 4 μg/mL.

### 3.2 RNA extraction, cDNA library construction, and RNA-seq

Following cell harvest, total RNA was extracted using TRIzol reagent (Invitrogen, Carlsbad, CA, USA) strictly according to the manufacturer’s instructions. RNA purity and concentration were initially quantified using a NanoDrop 2000 spectrophotometer (Thermo Fisher Scientific, USA). RNA integrity was subsequently evaluated via an Agilent 2100 Bioanalyzer (Agilent Technologies, Santa Clara, CA, USA), ensuring an RNA Integrity Number (RIN) ≥ 7.0 for all processed samples.

To preserve both coding and non-coding RNA profiles, ribosomal RNA (rRNA) was depleted from 1 μg of total RNA per sample using the Ribo-Zero Gold rRNA Removal Kit (Illumina, San Diego, CA, USA). The remaining RNA was chemically fragmented into short sequences using divalent cations under elevated temperature. Strand-specific cDNA libraries were constructed utilizing the NEBNext® Ultra™ Directional RNA Library Prep Kit for Illumina (Gibco, USA) following the instruction. Briefly, first-strand cDNA was synthesized using random hexamer primers and M-MuLV Reverse Transcriptase, followed by second-strand cDNA synthesis wherein dUTP was incorporated in lieu of dTTP to preserve transcript directionality. Following end repair, A-tailing, and adapter ligation, the uracil-containing second strand was degraded using USER enzyme. The libraries were then PCR-amplified and purified using AMPure XP beads (Beckman Coulter, Brea, CA, USA). Library quality was validated on the Agilent 2100 system, and precise quantification was performed using a Qubit 2.0 Fluorometer (Life Technologies). Ultimately, the validated libraries were pooled and sequenced on an Illumina Novaseq™ 6000 platform (PE 150 mode) by LC-Bio Technology Co., Ltd. (Hangzhou, China).

### 3.3 Quality control of raw data generated from RNA-seq

Upon acquisition of the raw paired-end reads in FASTQ format, initial quality assessment was performed using FastQC (v0.11.9). To ensure downstream analytical precision, rigorous data filtration was executed utilizing Cutadapt (v3.0). The filtering criteria systematically removed: (1) reads containing adapter contaminants, (2) reads harboring >10% ambiguous bases (N), and (3) low-quality reads where more than 50% of bases possessed a Phred quality score (*Q* score) < 20. Post-filtering statistics confirmed the generation of high-quality datasets (Table 2): the un-treated samples (C1-C5) yielded between 35,784,270 and 40,970,654 valid reads (valid data ratios: 95.72%-96.30%), whereas the melittin-treated samples (T1-T5) produced between 33,584,378 and 37,489,822 valid reads (valid data ratios: 95.96%-96.75%). Furthermore, all datasets exhibited exceptional base-calling accuracy, achieving a *Q*20 of 100.0% and a *Q*30 ≥ 99.08%. The GC content distribution ranged from 49.50% to 51.50%, which aligns with the theoretical GC content of the *Mus musculus* transcriptome.These stringent quality metrics verified the reliability of the sequencing data, providing a robust foundation for downstream differential gene expression, functional enrichment, and regulatory network analyses.

**Table 1.** Contents of the rawdata/raw/neg and rawdata/raw/pos folder.

| File Pattern | Count | Description |
| --- | --- | --- |
| Control_1.fastq.gz to<br>Control_5.fastq.gz | 5 | Raw data files for control group samples (Control.), with biological replicates numbered 1–5. |
| Control_1.fastq.gz.md5 to<br>Control_5.fastq.gz.md5 | 5 | MD5 checksum files for the corresponding control group .raw files, ensuring data integrity. |
| Treatment_1.fastq.gz to<br>Treatment_5.fastq.gz | 5 | Raw data files for melittin-treated group samples (Treatment), with biological replicates numbered 1–5. |
| Treatment_1.fastq.gz.md5 to<br>Treatment_5.fastq.gz.md5 | 5 | MD5 checksum files for the corresponding treatment group FASTQ files, ensuring data integrity. |

**Table 2.** Quality control of RNA-seq data.

| Sample | Raw data | Valid data | Valid ratio(reads) | $Q_{20}\%$ | $Q_{30}\%$ | GC content (%) |
| --- | --- | --- | --- | --- | --- | --- |
| C1 | 40,218,712 | 38,498,298 | 95.72 | 100.00 | 99.12 | 50.50 |
| C2 | 40,653,492 | 38,979,678 | 95.88 | 100.00 | 99.08 | 50.50 |
| C3 | 37,158,750 | 35,784,270 | 96.30 | 100.00 | 99.11 | 50.50 |
| C4 | 38,331,636 | 36,838,434 | 96.10 | 100.00 | 99.10 | 51.50 |
| C5 | 42,654,050 | 40,970,654 | 96.05 | 100.00 | 99.13 | 51.50 |
| T1 | 34,996,662 | 33,584,378 | 95.96 | 100.00 | 99.09 | 49.50 |
| T2 | 37,511,766 | 36,089,764 | 96.21 | 100.00 | 99.09 | 51.50 |
| T3 | 35,985,566 | 34,654,750 | 96.30 | 100.00 | 99.09 | 51.50 |
| T4 | 36,687,182 | 35,379,464 | 96.44 | 100.00 | 99.11 | 50.50 |
| T5 | 38,749,618 | 37,489,822 | 96.75 | 100.00 | 99.10 | 51.50 |

### 3.4 Read mapping and transcriptome assembly

Following quality control, the retained high-quality valid reads (Clean data) were aligned to the *Mus musculus* reference genome (Ensembl release 112) utilizing HISAT2 (v2.2.1) with default parameters. The resulting SAM files were converted to sorted BAM files using SAM tools (v1.10). The mapping results demonstrated high fidelity, with total mapping ratios ranging from 95.44% to 97.03% and uniquely mapped ratios between 88.86% and 89.97% across all samples (Table 3). Based on these high-confidence alignments, transcript reconstruction and the estimation of expression abundances were executed utilizing StringTie (v2.1.6) guided by the reference annotation.

**Table 3.**
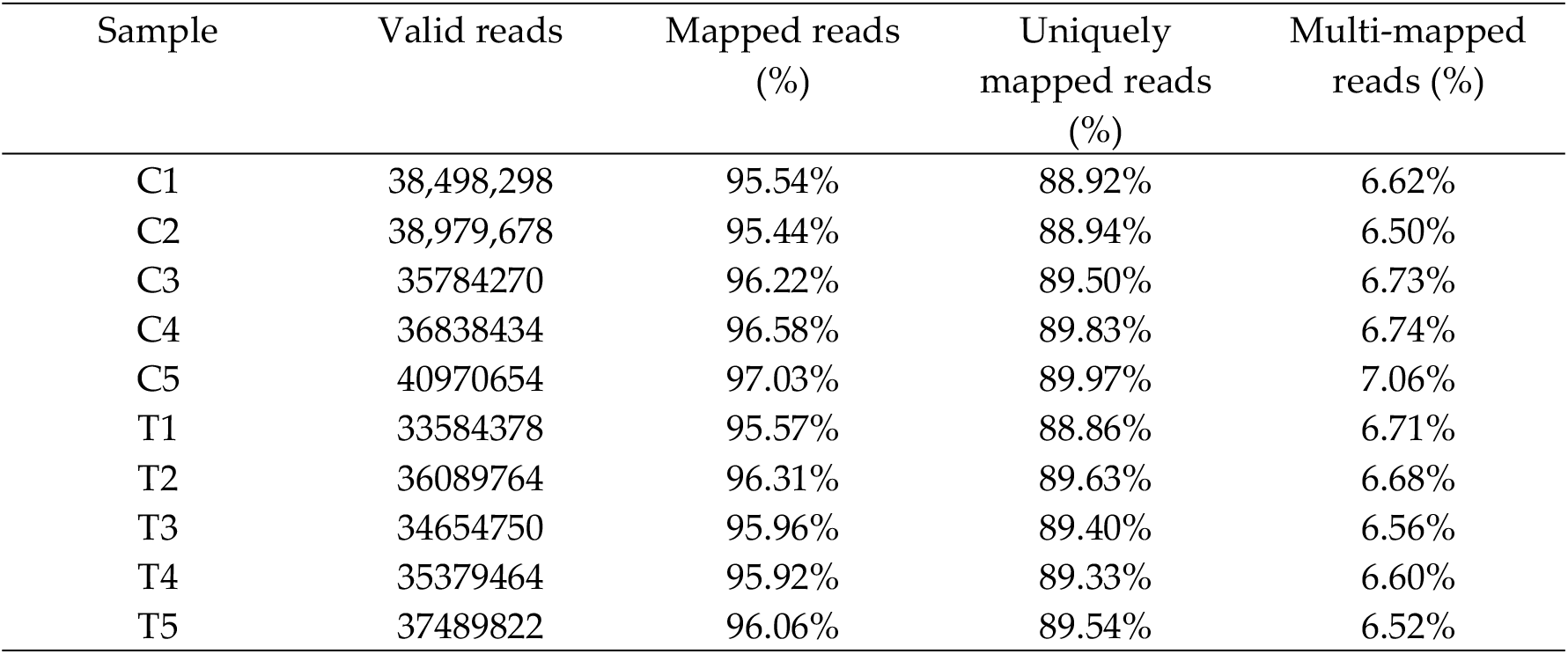
Overview of reads mapped to the *Mus musculus* reference genome.

| Sample | Valid reads | Mapped reads (%) | Uniquely mapped reads (%) | Multi-mapped reads (%) |
| --- | --- | --- | --- | --- |
| C1 | 38,498,298 | 95.54% | 88.92% | 6.62% |
| C2 | 38,979,678 | 95.44% | 88.94% | 6.50% |
| C3 | 35784270 | 96.22% | 89.50% | 6.73% |
| C4 | 36838434 | 96.58% | 89.83% | 6.74% |
| C5 | 40970654 | 97.03% | 89.97% | 7.06% |
| T1 | 33584378 | 95.57% | 88.86% | 6.71% |
| T2 | 36089764 | 96.31% | 89.63% | 6.68% |
| T3 | 34654750 | 95.96% | 89.40% | 6.56% |
| T4 | 35379464 | 95.92% | 89.33% | 6.60% |
| T5 | 37489822 | 96.06% | 89.54% | 6.52% |

### 3.5 Quantification of gene expression and differential analysis

Gene expression levels were quantified as Fragments Per Kilobase of transcript per Million mapped reads (FPKM) using StringTie, serving as a normalized metric for visualization and inter-sample comparison. To assess biological reproducibility and identify potential outliers, Principal Component Analysis (PCA) and Pearson correlation coefficient matrices were computed using the ggplot2 and pheatmap packages in R (v4.1.0), respectively.

For differential expression analysis, raw read counts for each gene were extracted using featureCounts (v2.0.1). Genes with low expression (total read counts < 10 across all samples) were pre-filtered to improve statistical power. The differential expression analysis between the melittin-treated and control cohorts was conducted utilizing the DESeq2 R package (v1.32.0). DESeq2 provides statistical routines for determining differential expression using a model based on the negative binomial distribution. To control the False Discovery Rate (FDR), raw *P* values were adjusted via the Benjamini-Hochberg (BH) procedure to generate *Q* values. Transcripts exhibiting an absolute log_2_(Fold change) ≥ 1 and an adjusted *Q* value < 0.05 were strictly defined as significantly DEGs, yielding a total of 158 up- and 96 down-regulated transcripts.

### 3.6 Functional enrichment analysis

To systematically unravel the biological implications and underlying molecular mechanisms of the identified DEGs, comprehensive functional enrichment analyses were executed utilizing the clusterProfiler R package (v4.0.0) (https://release.geneontology.org/2021-04-10/). Initially, the DEGs were mapped to specific terms within the GO database (v21.04.10), which categorizes gene functions into Biological Process (BP), Cellular Component (CC), and Molecular Function (MF) ontologies. Concurrently, pathway-level enrichment was performed utilizing the KEGG database (v21.04.10) (https://www.kegg.jp/kegg/pathway.html) to identify significantly perturbed signaling cascades and metabolic networks.

The statistical significance of both GO term and KEGG pathway enrichment was calculated utilizing a hypergeometric test, comparing the proportion of DEGs mapped to a specific term against the background distribution of all identified transcripts. A Benjamini-Hochberg adjusted *P* value (*Q* value) < 0.05 was employed as the threshold for significant enrichment. To quantitatively evaluate the degree of enrichment, a Rich Factor was calculated (defined as the ratio of DEGs mapped to a specific term/pathway to the total number of background genes annotated to that same pathway). The top 20 most significantly enriched terms and pathways were visualized using dot plots and bar charts generated via the enrichplot and ggplot2 R packages.

## Author Contributions

All the authors participated in the conception and design of the study. RH. Zhang, YW. Zhang obtained and analyzed the data. MY. Wang collected the samples. YL. Li, JR. Jiang organized the data and drafted the manuscript. DF. Chen, TZ. Yan, and R. Guo administrated the project, supervised the study, and revised the manuscript.

## Funding

This study was supported by the Foundation Research Project of Dongguan City (2251800400402), the Scientific and Technical Innovation Fund of Fujian Agriculture and Forestry University (KFb22060XA).

## Institutional Review Board Statement

Not applicable.

## Informed Consent Statement

Not applicable.

## Data Availability Statement

The raw transcriptome RNA sequencing data in FASTQ format, corresponding MD5 checksum files for data integrity validation, and processed gene expression matrices generated and analyzed in this study are openly available in the National Genomics Data Center (NGDC) Genome Sequence Archive. All raw data are available at the NGDC BioProject PRJCA068439 as described above, under the unique accession number PRJCA068439. No restricted or private data are included in this study.

## Acknowledgments

All of the authors appreciate the constructive and valuable comments from editors and reviewers.

## Conflicts of Interest

The authors declare no conflicts of interest.

The following abbreviations are used in this manuscript:

Abbreviation: Full Name in English
Akt: Protein Kinase B
Bax: Bcl-2 Associated X Protein
Bcl-2: B-cell Lymphoma 2
BH: Benjamini-Hochberg
BP: Biological Process
CC: Cellular Component
cDNA: complementary Deoxyribonucleic Acid
DEGs: Differentially Expressed Genes
DMEM: Dulbecco’s Modified Eagle Medium
dTTP: Deoxythymidine Triphosphate
dUTP: Deoxyuridine Triphosphate
FBS: Fetal Bovine Serum
FDR: False Discovery Rate
FPKM: Fragments Per Kilobase of transcript per Million mapped reads
GC: Guanine-Cytosine
GO: Gene Ontology
GSA: Genome Sequence Archive
KEGG: Kyoto Encyclopedia of Genes and Genomes
MD5: Message-Digest Algorithm 5
MF: Molecular Function
mRNA: messenger Ribonucleic Acid
NGDC: National Genomics Data Center
PBS: Phosphate Buffered Saline
PCA: Principal Component Analysis
PE150: Paired-End 150 base pairs
PI3K: Phosphatidylinositol 3-Kinase
*Q*20: Phred Quality Score 20
*Q*30: Phred Quality Score 30
RIN: RNA Integrity Number

## Disclaimer/Publisher’s Note

The statements, opinions and data contained in all publications are solely those of the individual author(s) and contributor(s) and not of MDPI and/or the editor(s). MDPI and/or the editor(s) disclaim responsibility for any injury to people or property resulting from any ideas, methods, instructions or products referred to in the content.

## Notes

https://ngdc.cncb.ac.cn

### Competing Interest Statement

The authors have declared no competing interest.

